# Evaluating the Phytotoxicity of Methanolic Extracts of *Parthenium hysterophorus* L. on Selected Crops and Weeds

**DOI:** 10.1101/2022.01.21.477310

**Authors:** H.M. Khairul Bashar, Abdul Shukor Juraimi, Muhammad Saiful Ahmad-Hamdani, Md Kamal Uddin, Norhayu Asib, Md. Parvez Anwar, Ferdoushi Rahaman, Mohammad Amdadul Haque, Akbar Hossain

**Author notes:** Allcorrespondence (A.S.J.); (H.M.K.B.).

## Abstract

Herbicides made from natural molecules may be a good environmentally friendly alternative to synthetic chemical herbicides for weed control. As a result, this investigation was carried out to ascertain the phytotoxicity of *Parthenium hysterophorus* L. as well as to identify its phenolic components. Germination of seeds and development of seedlings of *Vigna subterranea* (L.) Verdc, *Raphanus sativus* (L.) Domin, *Cucurbita maxima* Duchesne., *Cucumis sativus* L., *Solanum lycopersicum* L., *Capsicum frutescens* L., *Zea mays* L., *Abelmoschus esculentus* (L.) Moench, *Daucus carota* L., *Digitaria sanguinalis* (L.) Scop and *Eleusine indica* (L.) Gaertn were investigated using *P. hysterophorus* leaf, stem, and flower methanol extracts. Six concentrations (25, 50, 75, 100, and 150 g L^−1^) were comparison to the control (distilled water). The concentration of extracts increased, the rate of the seed sprouting and seedling growth decreased. EC_50_ values showed that the extraction of leaf of *P. hysterophorus* (811) was phytotoxic in comparison to the stem (1554) and flower (1109) extract. According to PCA analysis, *Raphanus sativus, Solanum lycopersicum, Capsicum frutescens, Abelmoschus esculentus, Daucus carota, Digitaria sanguinalis*, and *Eleusine indica* were all very susceptible to allelochemicals. A LC-MS analysis revealed that the *P. hysterophorus* leaf extract contained 7 phenolic compounds that were responsible for inhibition. These studies also revealed that the leaf of *P. hysterophorus* is a major source of phytotoxicity, which could be valuable in the future for developing a natural herbicide.

## 1. Introduction

Parthenium (*Parthenium hysterophorus* L.) is a noxious herb that has now invaded 46 countries and extended its spread from a few islands to around the world, there have been eleven minor and eight major introductions [1]. Its high invasiveness is associated with several factors, including a higher number of seeds production, highly competitive and rapidly expanding, biological plasticity of the life cycle, allelopathic ability and high survival ability against biotic and abiotic stresses [2–5].

Allelopathy is described as chemical’s positive or negative impacts substances formed primarily by plant, microbe, and fungal secondary metabolism on the growth and establishment of neighbouring plants or microorganisms, as well as the dynamical processes of agricultural and natural eco-systems [6]. It’s a complicated phenomenon that’s influenced by a variety of internal and external circumstances. Due to its intricacy, the explanation is a difficult endeavour that necessitates knowledge from a variety of professions [7]. Allelochemicals are plants that release secondary metabolites into the environment. They are anti-inflammatory substances that belong to a variety of chemical classes, primarily phenolic compounds and terpenoids [8]. Bhadoria [9] provided a more comprehensive summary of allelochemicals’ that affects plant growth and development. All plant organs (stems, leaves, rhizomes, roots, flowers, pollen, fruits, seeds) contain allelochemicals, which are released through volatilization, leaf leaching, plant material breakdown, and root exudation. In some way, membrane stability, cell division, elongation, shape and permeability, enzyme activity, and respiration of plants are all influenced. Photosynthesis, protein synthesis, nucleic acid metabolism, and other direct and indirect ways of action cause seed sprouting suppression and limited seedling development [8]. In addition, the microbial breakdown of soil allelochemicals has an impact on the effective dose of allelochemicals that can inhibit plants [10,11].

Herbicides have been the least expensive and principal method of weed control in developing countries for about 50 years [12]. Herbicides, on the other hand, pose significant risks to agriculture, human health, and the environment. However, increasing crop production without using chemical herbicide is an urgent challenge in crop production. Manual weed management is the most effective and long-term solution for weed management. So, accurate weed control is necessary for food security throughout the world. Therefore, researchers are motivated to seek alternatives because of the labour movement from agriculture to others, and weed biotypes resistant to traditional synthetic pesticides [13]. This strategy will aid in reducing reliance on chemical herbicides, reducing the likelihood of weed resistance to herbicides, reducing health risks and environmental damage, and strengthening the national economy. In the meantime, there are a variety of possible allelochemicals in aerial sections (e.g. leaves) of *Parthenium* weed have been confirmed by several earlier studies; among them p-anisic acid (C_8_H_8_O_3_), p-coumaric acid (C_9_H_8_O_3_), caffeic acid (C_9_H_8_O_4_), ferulic acid (C_4_H_4_O_4_), fumaric acid (C_4_H_4_O_4_), p-hydroxybenzoic acid (C_7_H_6_O_3_), neochlorogenic acid (C_16_H_18_O_9_), protocatechuic acid (C_7_H_6_O_4_), aerulic acid, chlorogenic acid (C_16_H_18_O_9_) and vanillic acid (C_4_H_4_O_4_) are the most important [14,15] sprouting and development of a plant species in abundance, natural plants are included and different crops and pasture species can be inhibited by these chemicals [16]. Wheat, maize, and horse gram [5], lentil [17,18] and other field crops. Hassan *et al*., [19] showed an inhibitory impact when exposed to parthenium extract. Dhawan and Gupta [15] reported that the extraction of diverse active phytochemicals with flavonoid concentrations works best using methanol as an extraction solvent.

However, there is insufficient evidence on the influence of Parthenium methanolic extracts on the sprouting and seedlings development of several crops especially Bambara groundnut weeds. The Bambara groundnut is a new crop for Malaysia, but there is information lacking on the suppression of allelopathy on Bambara groundnut weeds by different parts of *P. hysterophorus*. The current study aimed to find out the allelopathic capacity of Parthenium in a laboratory experiment to evaluate the allelopathic suppression of weeds by *P. hysterophorus* in Bambara groundnut weeds. The research was directed with the following objectives (1) to evaluate the phytotoxicity of methanol extracts made from the aerial portions of *P. hysterophorus* on target species in order to develop bioherbicides based on natural products (2) LC-MS was used to identify its phenolic derivatives.

## 2. Materials and Methods

### 2.1. Experimental location

Growth chamber research was carried out at Weed Science Lab in the Crop Science Department, Faculty of Agriculture, Universiti Putra Malaysia (3°02’ N, 101°42’ E, elevation 31 m), Malaysia. Temperature in the growth chapter was maintained at 25°C in throughout the experimental period.

### 2.2. Experimental Treatments and Design

Leaf, stem, and floral parts of parthenium was applied at different concentration viz., 0, 25, 50, 75, 100, and 150 g L^−1^ [42]. All treatments were arranged in a completely randomized design (CRD) and repeated four times.

### 2.3. Plant Materials and Preparation of Seeds

For extraction of the leaf of *P. hysterophorus* plants, plant materials were taken from Ladang Infoternak farm in Sungai Siput, Perak, Malaysia, and also grown in the net house of field 15 at University of Putra Malaysia, Selangor, Malaysia. The above-ground part of the plants was collected just before maturity, rinsed several times using tap water to eliminate dust elements, then air-dried at ambient temperature (24-26°C) for three weeks. The leaves, stems, and flowers were divided and bulked up into three main parts. In a laboratory blender, both bulked plant components were ground into fine dust and sieved through a 40-mesh sieve.

The inhibitory action of *P. hysterophorus* was investigated on nine plant species. Bambara groundnut (*Vigna subterranea* L. Verdc), radish (*Raphanus sativus* L. Domin), sweet gourd (*Cucurbita maxima* Duchesne), tomato (*Solanum lycopersicum* L.), cucumber (*Cucumis sativus* L.), chili (*Capsicum frutescens* L.), corn (*Zea mays* L.), carrot (*Daucus carota* L.) and okra (*Abelmoschus esculentus* L. Moench) and two weed species goosegrass (*Eleusine indica* L. Gaertn)] and [crab grass (*Digitaria sanguinalis* L. Scop). Crop seeds were attained from Sin Seng Huat Seeds Sdn Bhd company in Malaysia, while seeds of grasses were personally picked from the Universiti Putra Malaysia’s agricultural field. The seeds were cleaned, air-dried, and stored in airtight containers maintain at −18° C. The vegetable crops are chosen for the determination of ecological effects of allelopathic substances as they represented commonly used species in the field that’s are recommended by US EPA [54]. They belong to different plant families and can provide great genetic diversity. The seeds germinated 86-95% of the time, according to a random test.

### 2.4. Extract Preparation

The extracts were made according to the procedure published by [55] and [42]. Accurately 100 g powder from leaves, stems, and flowers of parthenium were placed in a conical flask and allowed to soak in 1L of 80% (v/v) methanol separately. After that, the conical flask was wrapped in paraffin and shaken for 48 hours at 24-26°C room temperatures in an Orbital shaker at 150 rpm agitation speed. To remove debris, cheesecloth in four layers were used to filter the mixtures and centrifuged for one hour at 3000 rpm in a centrifuge (5804/5804 R, Eppendorf, Germany). A single layer of Whatman No. 42 filter paper was used to filter the supernatant. A 0.2-mm Nalgene filter was used to filter the solutions once more to avoid microbial development (Lincoln Park, NJ-based Becton Dickinson percent Labware). Using a rotary evaporator (R 124, Buchi Rotary Evaporator, Germany), the solvents were evaporated from the extract to dryness (a thick mass of coagulated liquid) under vacuum at 40° C and the sample was then collected. From a 100 g sample of *P. hysterophorus* powder, the average extracted sample was 17.56 g.

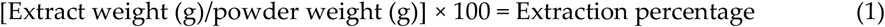

For the bioassay, each stock extract from *P. hysterophorus* leaves, stems, and flowers were diluted in sterile distilled water to provide extract concentrations of 25, 50, 75, 100, and 150 g L^−1^, while purified water was served as control. All extracts were stored at 4° C in the dark until use.

For LC-MS analysis, 100% HPLC GRADE methanol (20 mL) was diluted with the crude sample (20 mg) and filtered through 15-mm, 0.2-μm syringe filters (Phenex, Non-sterile, Luer/Slip, LT Resources Malaysia).

### 2.5. Germination and growth bioassays

Healthy, uniform seeds were gathered and treated with 0.2% potassium nitrate for 24 hours (KNO_3_) before being rinsed with distilled water. Twenty Bambara groundnut and sweet gourd seeds and thirty seeds of radish, cucumber, tomato, chili, corn, okra, carrot, crabgrass, and goosegrass were set up in a sterilized Petri dish with Whatman No. 1 filter paper (90 × 15 mm). 10 mL of extract of each concentration (25, 50, 75, 100, and 150 g L^−1^) was delivered in Petri dishes, distilled water serving as a control. In a growth chamber, all Petri dishes were inserted. and incubated at 30° C/20°C (day/night) temperature under fluorescent light (8500 lux) on photoperiod 12 h day/12 h night maintained 30-50% relative humidity. To facilitate gas exchange, the petri dish lids were not sealed.

### 2.6. Identification of phenolic derivatives in P. hysterophorus leaves, stems, and flowers extracted in methanol

The LC-MS was used to identify the chemical contents of the extracts. The phytochemical compounds of the methanol extracts were performed using LC-MS followed by [56]. LC-MS analysis was performed using Agilent spectrometry equipped with a binary pump. The LC-MS was interfaced Agilent 1290 Infinity LC system coupled to Agilent 6520 accurate-mass Q-TOF mass spectrometer with a dual ESI source. Full-scan mode from m/z 50 to 500 was performed with a source temperature of 125°C. The column of Agilent zorbax eclipse XDB-C18, narrow-bore 2.1×150 mm, 3.5 microns (P/N: 930990-902) was used with the temperature 30°C for the analysis. A- 0.1% formic acid in water and B −0.1% formic acid in methanol were used as solvents. Isocratic elution was used to supply solvents at a total flow rate of 0.1 mL minutes^−1^. MS spectra were collected in both positive and negative ion modes. The drying gas was 300° C, with a 10L min-1 gas flow rate and a 45-psi nebulizing pressure. Before analysis, 1 ml of concentration. sample extracts were diluted with methanol and filtered through a 0.22 m nylon filter. The extracts were injected into the analytical column in 1 μl volume for analysis. The mass fragmentations were discovered using an Agilent mass hunter qualitative analysis B.07.00 (Metabolom-ics-2019.m) tool and a spectrum database for organic chemicals.

### 2.7. Data collection

The germination percentage, radicle, and hypocotyl length were measured with a ruler at seven days after seeding. The radicle and hypocotyl length was assessed by software Image J [57] while the inhibition (%) of *P. hysterophorus* extracts on a radicle, and hypocotyl length was computed following the formula used by Kordali [58]:

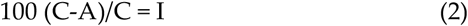

Here, “I” is the percentage of inhibition, “C” control’s mean growth and development and “A” is the aqueous extracts’ mean growth and development.

### 2.8. Statistical Analysis

On pooled (two seasons) data, a one-way analysis of variance (ANOVA) was used to regulate any significant variances among concentrations and control. To calculate the difference between the concentration means, the Tukey test (SAS 9.4) with a 0.05 probability level was utilised. EC_r50_, EC_g50_, and EC_h50_ were used to compute real dosages accomplished of suppressing 50% of germination, radicle development, and hypocotyl growth. Based on the suppression of germination (percentage), radicle, and hypocotyl development, Probit analysis was used to compute the EC_g50_, EC_r50_, and EC_h50_ values. From each tested plant, a rank was determined by using the following equation to calculate an index (Re) for each of the most active extracts and plants that are the most susceptible:

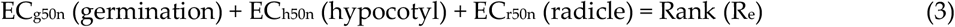

Where Re is the plant’s rank n, EC_r50n_, EC_h50n_ and EC_g50n_ are the amounts of plant extract n that inhibit 50% germination, radicle, and hypocotyl length, respectively. The lowest Re value had the maximum active tissue extracts and the utmost sensitive plants, while the highest Re value had the least allelopathic effect of the extract.

The most common application of NTSYSpc 2.02e (Numerical Taxonomy and Multivariate Analysis System) is to do various types of agglomerative cluster analysis of some type of similarity or dissimilarity matrix and the quantity of extract sensitivity among the plants under investigation [59,60]. The principal component analysis (PCA) was used to re-validate Johnson’s cluster analysis [20].

## 3. Results

### 3.1. Inhibitory influence of P. hysterophorus on crop species

Different concentrations of methanolic extracts to the control, Parthenium leaf, stem, and flower concentrations and different crops had a significant influence on germination of seed, radicle, and hypocotyl length of the examined plants, as well as a rise in extract concentration. Parthenium extracts had a bit stimulatory impact on seed germination at 25 g L^−1^, but an inhibitory effect was observed at higher dosages (Figure 1).

**Figure 1.**
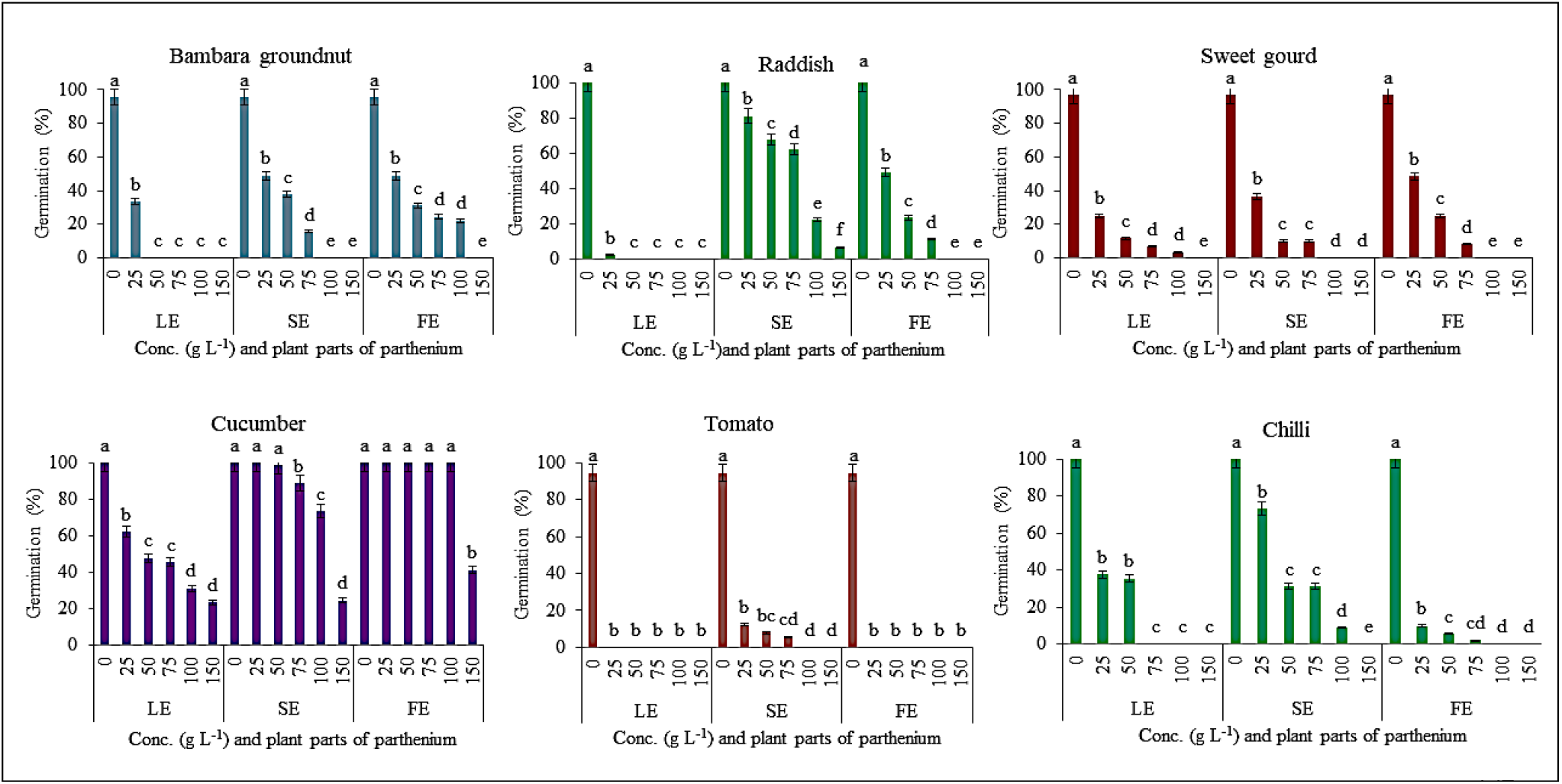
Showing the effect on germination% from different concentration levels of Parthenium aerial plant parts on crops (Bambara groundnut, raddish, sweet gourd, cucumber, tomato and chilli). Note: LE-Leaf extract, SE – Stem extract, FE – Flower extract.

Methanolic extract of leaf at 25 g L^−1^ significantly decreased the sprouting of all plants except sweet gourd, cucumber, and maize (p≤0.05), while, seed germination failure was seen in tomato, carrot, and goosegrass if the concentration level further increased. The maximum concentration resulted in 100% germination failure in all crops except cucumber (76%) and corn (65%) (Figure 1 & 2).

**Figure 2.**
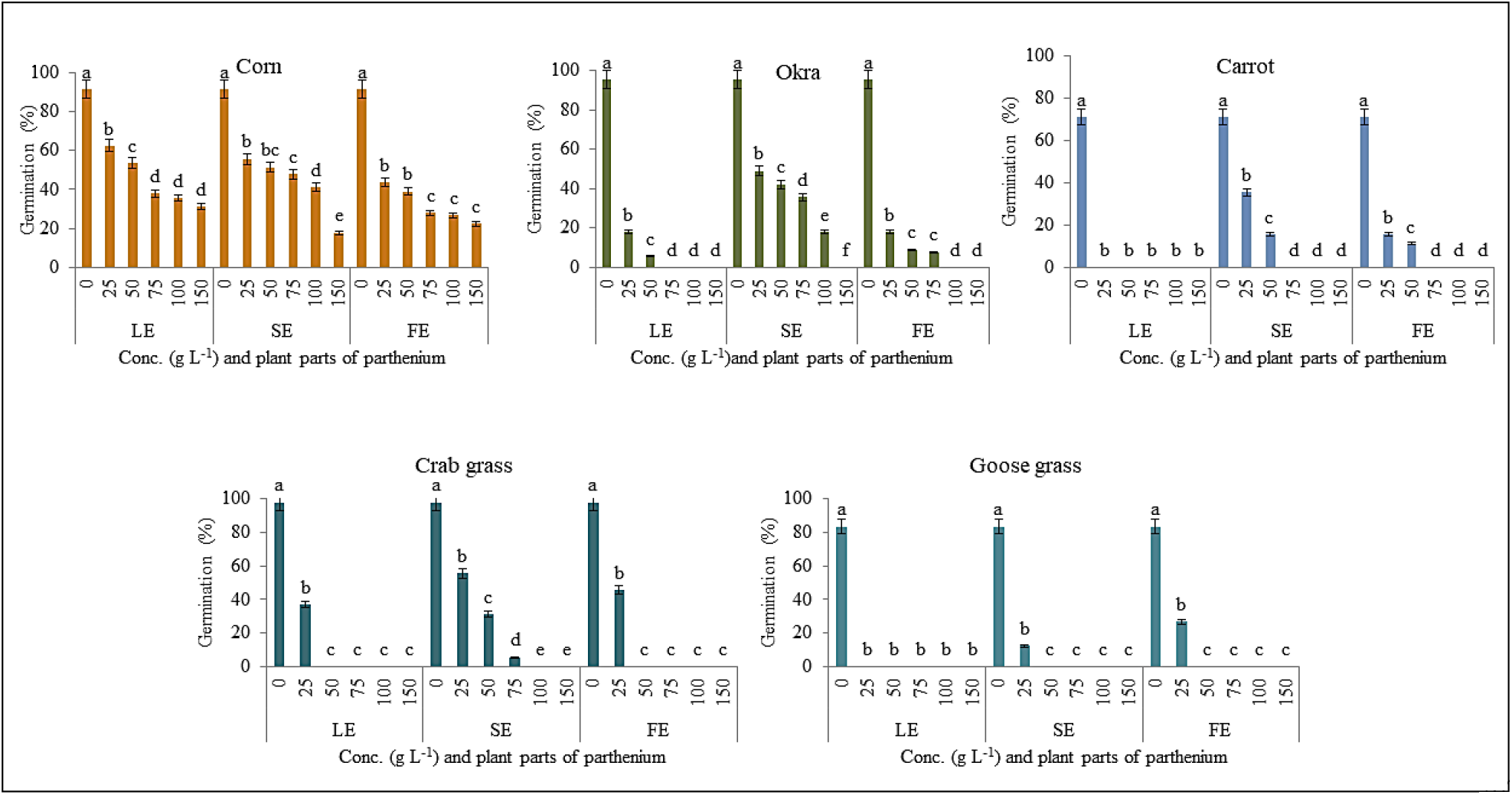
Showing the effect on germination% from different concentration levels of Parthenium aerial plant parts on crops (corn, okra and carrot) and weeds species (crab grass and goose grass). Note: LE-Leaf extract, SE – Stem extract, FE – Flower extract.

When the *P. hysterophorus* stem and flower extract were applied at lower doses (25, 50, and 75 g L^−1^), there was no significant reduction in germination (%). When the concentration was raised from 100 to 150 g L^−1^, the sprouting was substantially decreased between 1-100 percent in the stem and 48-100 percent in the flower extract among the indicator plants, while it was 61-100 percent in the leaf extract (Figure 1& 2). Among them, leaf extract was affected in many crops than stem and flower extract. On the other hand, germination (%), radicle, and hypocotyl length were significantly decreased at 50 to 100 g L^−1^ leaf extracts (Table 1). Increasing the concentration level eventually reduced the germination percentage over time. Both extracts of Parthenium inhibit the germination percentage of examined indicator both weed species (Figure 2).

**Table 1.**
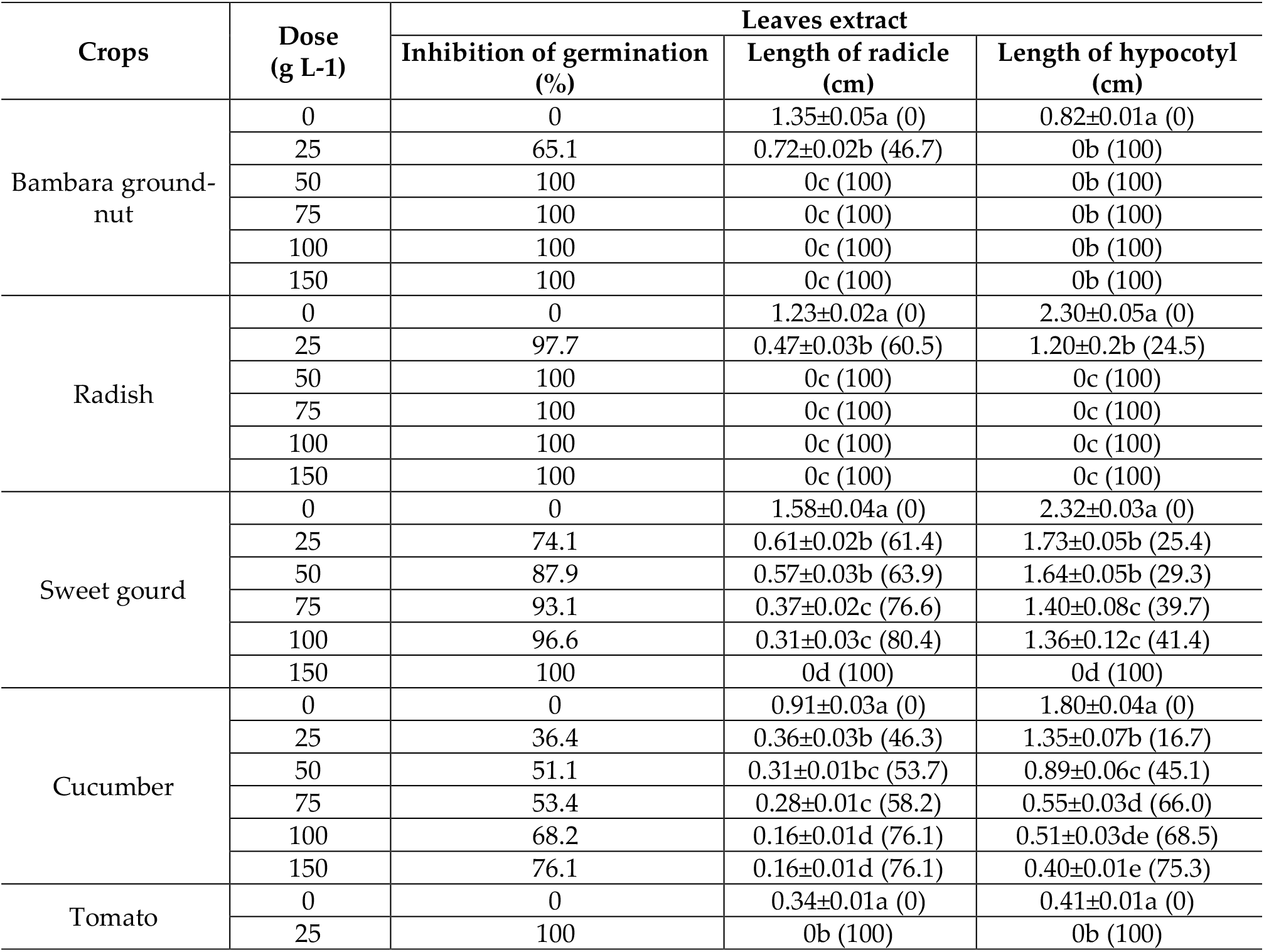
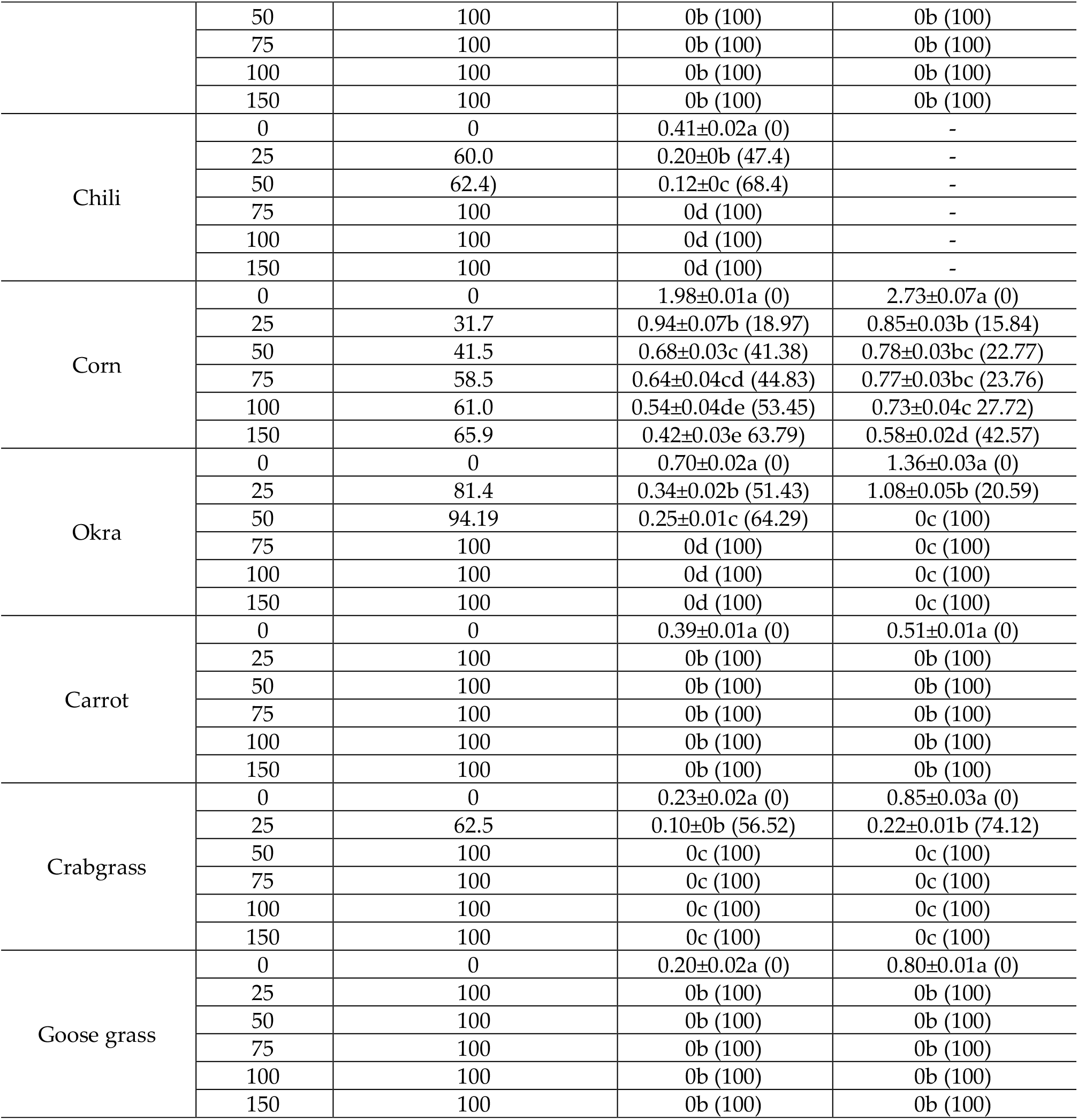
Effect of leaves extracts of *Parthenium hysterophorus* with Methanol on germination, radicle and hypocotyl length, and percent (%) inhibition of different crops.

Methanol extracts had a significant phytotoxic influence on all the crops on radicle and hypocotyl length studied at varied doses except sweet gourd, cucumber, and corn. Extraction of the leaf at doses more than or equal to 50 g L^−1^ substantially decreased the radicle length of target plants (p≤0.05) (Table 1). With 100 to 150 g L^−1^ stem and flower extract, root development of certain plants was decreased by more than half, whereas the uppermost concentration of the leaf extract (100 to 150 g L^−1^) resulted in no root development such as Bambara groundnut, radish, chili, okra, etc (Table 1). From the concentration level of 100 to 150 g L^−1^ Parthenium extract radicle length showed the inhibition level 53-100%, 36-100%, and 10-100% were from leaf, stem, and flower, respectively (Table 1, 2 & 3). As a result, the leaf extract had a higher concentration than the others (Table 1).

**Table 2.**
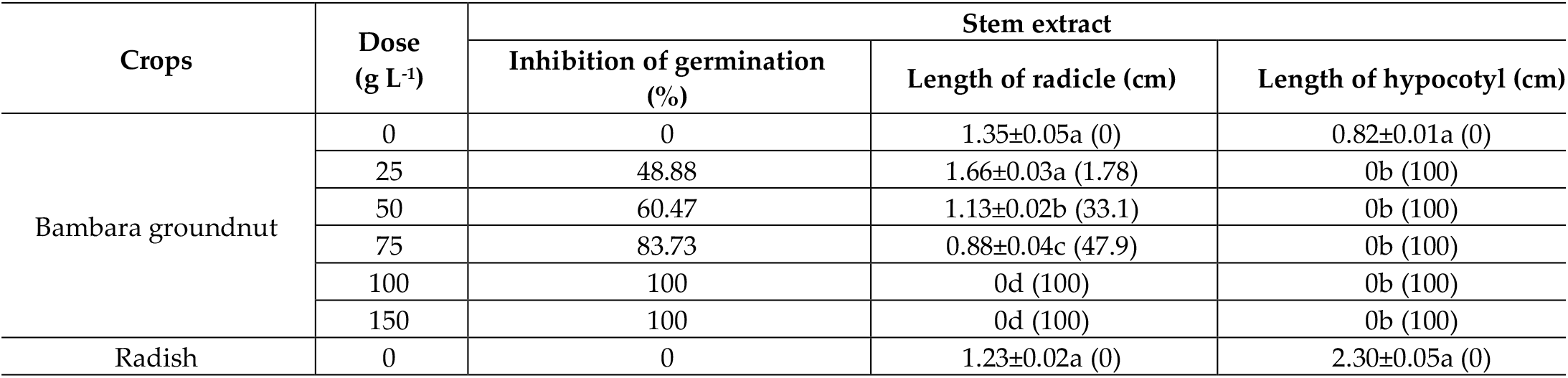
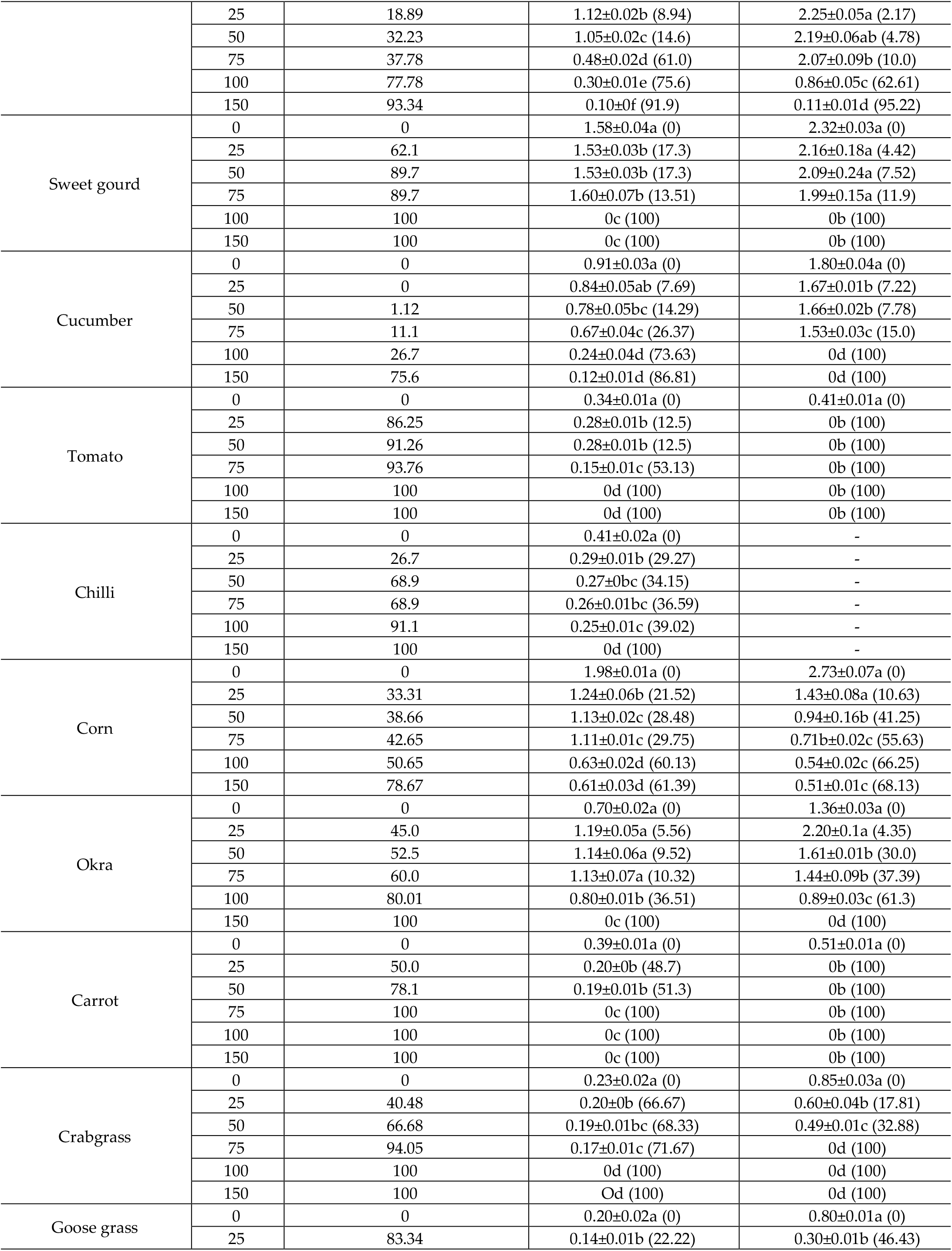
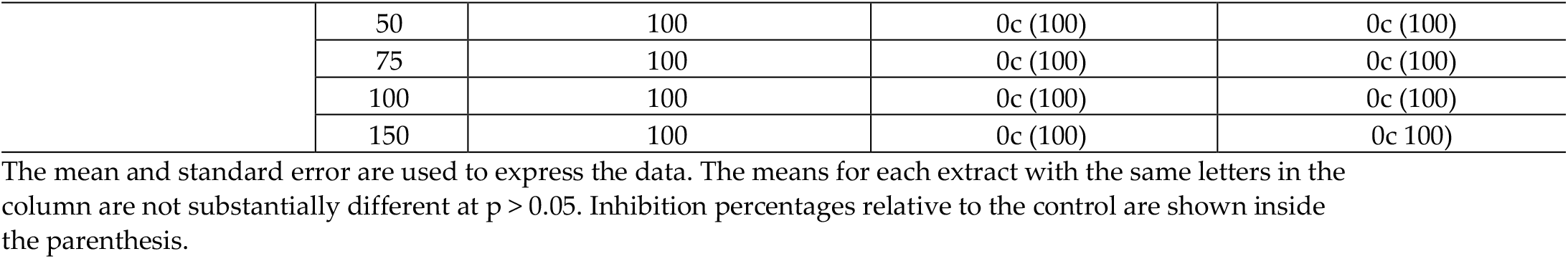
Effect of stem extracts of *Parthenium hysterophorus* with Methanol on germination, radicle and hypocotyl length, and percent (%) inhibition of different crops.

**Table 3.**
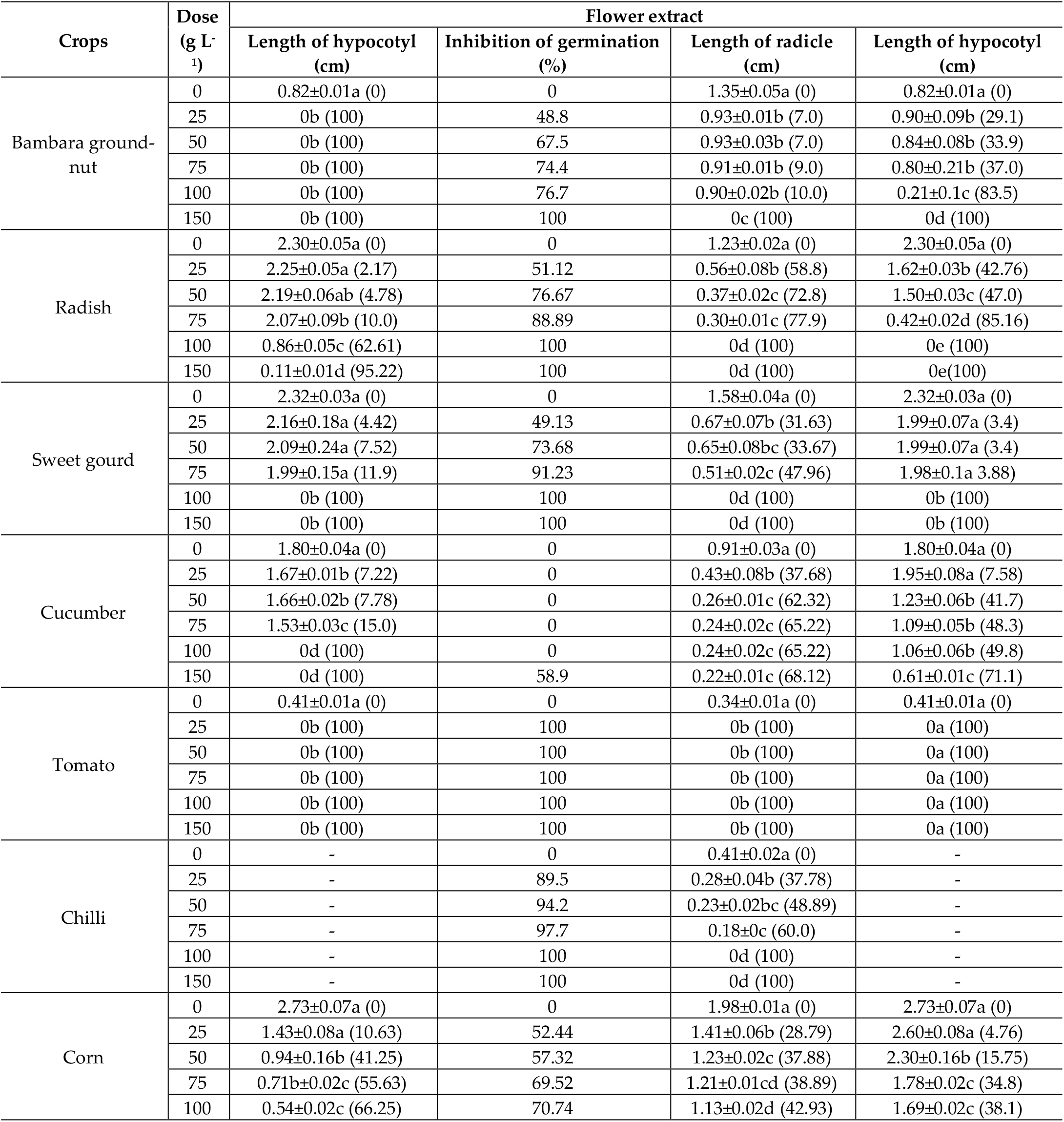
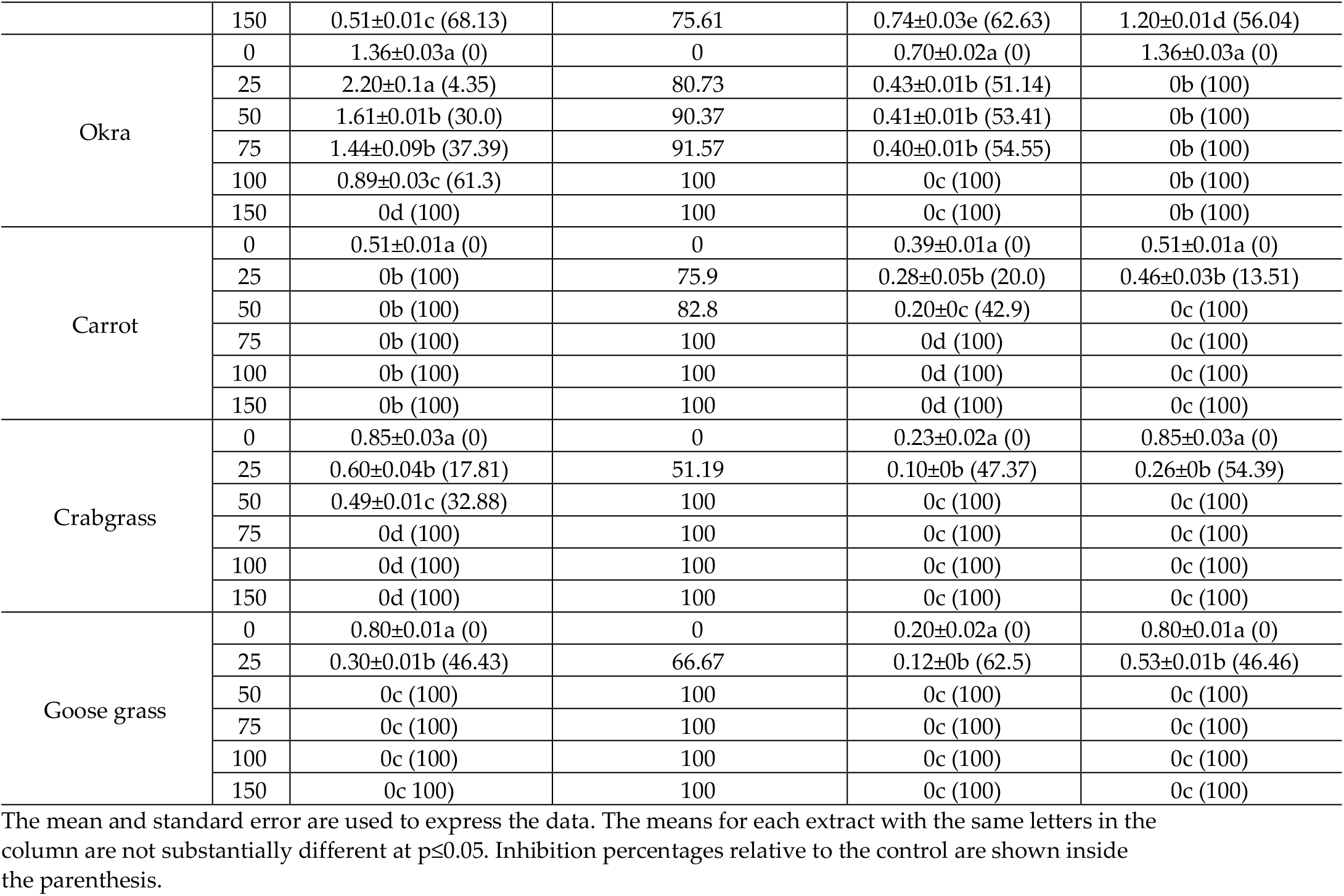
Effect of flower extracts of *Parthenium hysterophorus* with Methanol on germination, radicle and hypocotyl length, and percent (%) inhibition of different crops.

Furthermore, we observed that the weed crabgrass and goosegrass were severely affected by leaf and flower methanol extract in the doses of 50 to 150 g L^−1^, but stem extract was affected by doses of 100 to 150 g L^−1^. So, it was observed that severely affected weed by leaf than flower and stem plant parts. The amount of inhibition rose when the concentration level was raised. Different components of Parthenium reduced the shoot length of all examined plants by 27-100%, 61-100%, and 38-100%, respectively, at the doses of 100 to 150 g L^−1^.

### 3.2. The half inhibitory effect of Parthenium methanol extracts

Table 4 showed the half inhibitory (EC_50_) impact of Parthenium plant parts with methanol extracts, as well as the sensitivity of the evaluated starting growth parameters and plants. The efficacy of stem extract (1554) was lower than that of leaf extract (811), and it was followed by flower extract (1109) in all tested crops. The EC_50_ value showed some differences in sensitivity between the tested plant’s responses to the inhibitory influence of *P. hysterophorus* (Table 4). In case of leaf methanol extract corn, Cucumber, and sweet gourd were only impacted at higher concentrations. The rank value of these crops is 463, 144, and 108 respectively, it means that these crops are more tolerant, which shows that more doses need to destroy these plants. On the other hand, Bambara groundnut, radish, tomato, carrot, crabgrass, and goosegrass are more sensitive to leaf methanol extract next to chili (53) and okra (41).

**Table 4.** For the examined species, the rank value (Re) of *P. hysterophorus* methanol extract

| Target plants | Leaf extract |
| --- | --- |

|  | EC<sub>g50</sub> | EC<sub>r50</sub> | EC<sub>h50</sub> | Rank |
| --- | --- | --- | --- | --- |
|  | Values in g L<sup>-1</sup> |  |  |  |
| Bambara groundnut | 0 | 0 | 0 | 0 |
| Radish | 0 | 0 | 0 | 0 |
| Sweet gourd | 12.98 | 22.35 | 73.26 | 108.59 |
| Cucumber | 48.97 | 35.11 | 60.07 | 144.15 |
| Tomato | 0 | 0 | 0 | 0 |
| Chilli | 25.17 | 28.76 | 0 | 53.93 |
| Corn | 62.33 | 85.60 | 315.38 | 463.31 |
| Okra | 13.76 | 28.01 | 0 | 41.77 |
| Carrot | 0 | 0 | 0 | 0 |
| Crabgrass | 0 | 0 | 0 | 0 |
| Goosegrass | 0 | 0 | 0 | 0 |
| Rank | 163.21 | 199.83 | 448.71 | 811.75 |
|  | Stem extract |  |  |  |
| Bambara groundnut | 30.48 | 62.42 | 0 | 92.9 |
| Radish | 64.67 | 68.91 | 93.78 | 227.36 |
| Sweet gourd | 19.78 | 68.16 | 77.20 | 165.14 |
| Cucumber | 119.62 | 83.05 | 74.86 | 277.53 |
| Tomato | 7.09 | 61.23 | 0 | 68.32 |
| Chilli | 39.82 | 72.20 | 0 | 112.02 |
| Corn | 71.28 | 100.97 | 72.39 | 244.64 |
| Okra | 38.10 | 99.18 | 74.53 | 211.81 |
| Carrot | 26.59 | 31.04 | 0 | 57.63 |
| Crabgrass | 31.62 | 20.79 | 44.41 | 96.82 |
| Goosegrass | 0 | 0 | 0 | 0 |
| Rank | 449.05 | 667.95 | 437.17 | 1554.17 |
|  | Flower extract |  |  |  |
| Bambara groundnut | 29.20 | 114.79 | 56.50 | 200.49 |
| Radish | 26.33 | 23.86 | 35.43 | 85.62 |
| Sweet gourd | 27.54 | 49.26 | 75.02 | 151.82 |
| Cucumber | 143.50 | 37.19 | 83.40 | 264.09 |
| Tomato | 0 | 0 | 0 | 0 |
| Chilli | 6.46 | 40.59 | 0 | 47.05 |
| Corn | 23.31 | 108.16 | 126.93 | 258.4 |
| Okra | 10.14 | 34.24 | 0 | 44.38 |
| Carrot | 16.04 (0) | 41.69 | 0 | 57.73 |
| Crabgrass | 0 | 0 | 0 | 0 |
| Goosegrass | 0 | 0 | 0 | 0 |
| Rank | 282.52 | 449.78 | 377.28 | 1109.58 |

The quantities of extracts that inhibit 50% of germination, root, and hypocotyl, respectively, are designated as EC<sub>g50</sub>, EC<sub>r50</sub>, and EC<sub>h50</sub>.

Again, in case of stem methanol extract cucumber (277), corn (244), radish (227), okra (211), sweet gourd (165), and chili (112) are more tolerant and other crops are more sensitive. It was inhibited by the extract. On the contrary, in case of flower extract cucumber (264), corn (258), Bambara groundnut (200), sweet gourd (151) is more tolerant but tomato, crabgrass, and goosegrass with other crops are more sensitive. These findings revealed that *P. hysterophorys* leaf extract had a greater effect on plant development than flower and stem extract at all dosages.

Again, germination was seriously affected (163, 449, and 282) among the leaf, stem, and flower extracts indices, while radicle length (199, 667, and 449) and hypocotyl length (448, 437, and 377) were less affected to both plant sections. Overall, the methanol leaf extract of *P. hysterophorus* was very hazardous to all plants examined, particularly to germination, which was hindered at the lowest dosage.

### 3.3. Cluster and Principal Component Analysis (PCA)

Cluster analysis was also used to categorize distinct groups of plants with comparable responses to the inhibition of leaf, stem, and flower extracts by combining all three characteristics examined. Cluster analysis produced a dendrogram that revealed variation in sensitivity among the plants (Figure 3).

**Figure 3:**
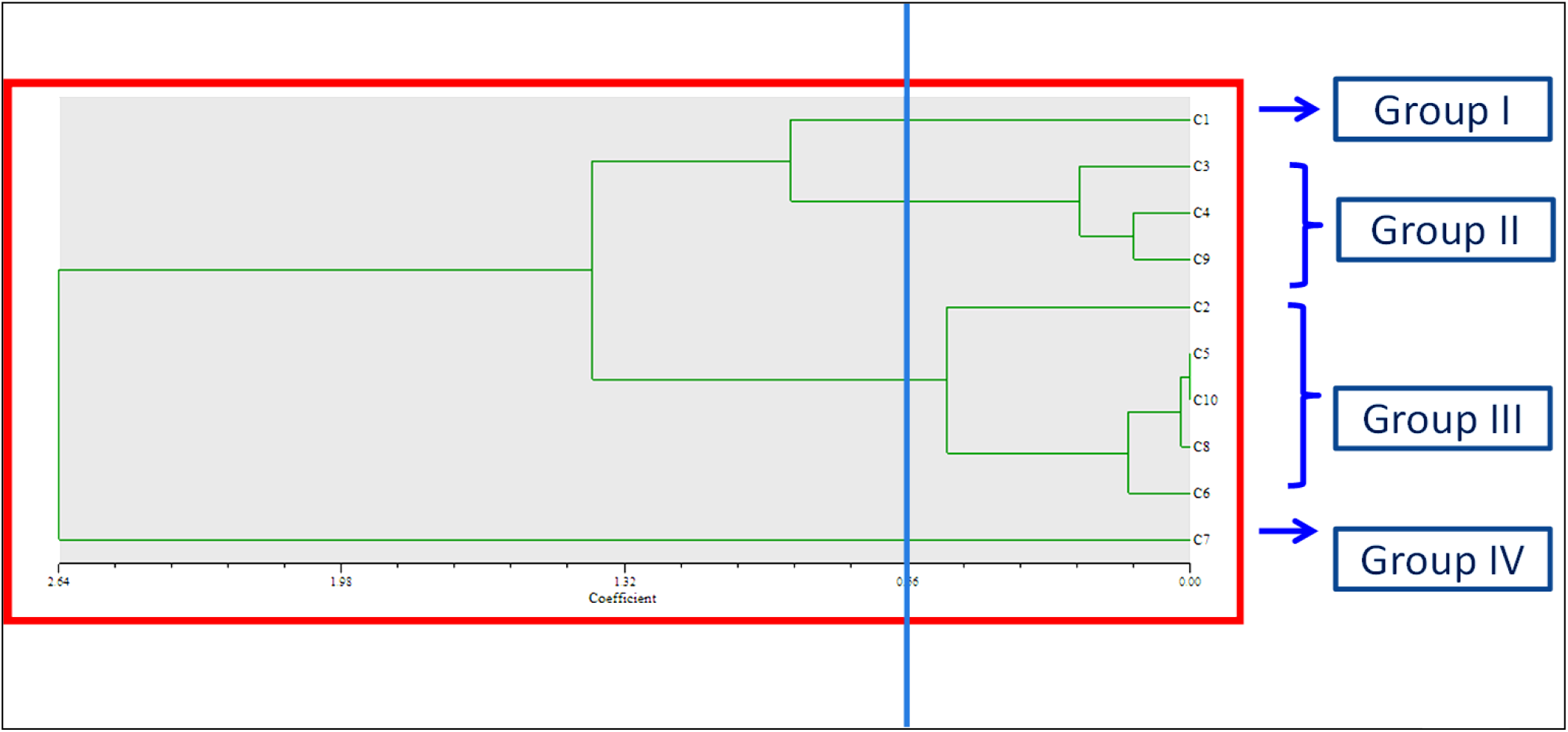
All indicator plants’ mean EC_50_ values for seed germination, radicle, and hypocotyl length are represented in a dendrogram (C_1_-Bambara groundnut, C_2_-Radish, C_3_-Sweet gourd, C_4_- Cucumber, C_5_- Tomato, C_6_- Chili, C_7_- Corn, C_8_- Okra, C_9_- Carrot, C_10_- Crabgrass, C_11_- Goosegrass) treated with the leaf, stem and flower extracts of *P. hysterophorus* with methanol revealed by non-overlapping (SAHN) UPGMA Method.

Plants may be divided into four classes based on how they react to leaf, stem, and flower extracts (Table 5). According to Table 5, group IV comprises tolerant monocot plants, whereas the dicot plants examined referred to the sensitive groups. Corn was recorded tolerant, whereas the moderately sweet gourd, cucumber, and carrot had an intermediate reaction to the phytotoxicity. The most vulnerable plants, on the other side, were Bambara groundnut, radish, tomato, crabgrass, goose grass, okra, and chili. Overall, the dicot plants were shown to be more active against the Parthenium extract than the monocots.

**Table 5.** Showing the similarity among the indicator plants.

| Clustering | Code | Name of the crop |
| --- | --- | --- |
| Group I | C<sub>1</sub> | Bambara groundnut |
| Group II | C<sub>3</sub>, C<sub>4</sub>, C<sub>9</sub> | Sweet gourd, Cucumber, Carrot |
| Group III | C<sub>2</sub>, C<sub>5</sub>, C<sub>10</sub>, C<sub>11</sub>, C<sub>8</sub>, C<sub>6</sub> | Radish, Tomato, Crabgrass, Goosegrass, Okra, Chili |
| Group IV | C<sub>7</sub> | Corn |

The principal component analysis (PCA), on the other hand, is a re-validation tool for cluster analysis. Johnson uses PCA to estimate the total variation that exists in a set of characters [20]. As shown by the eigenvector in the two-dimensional (Figure 4) and three-dimensional (Figure 5) graphical elucidations, the majority of the indicator plants were spread at short distances, while just two were dispersed at long distances. Bambara groundnut and Corn were the accessions that were farthest from the centroid, whilst other accessions were close to it.

**Figure 4:**
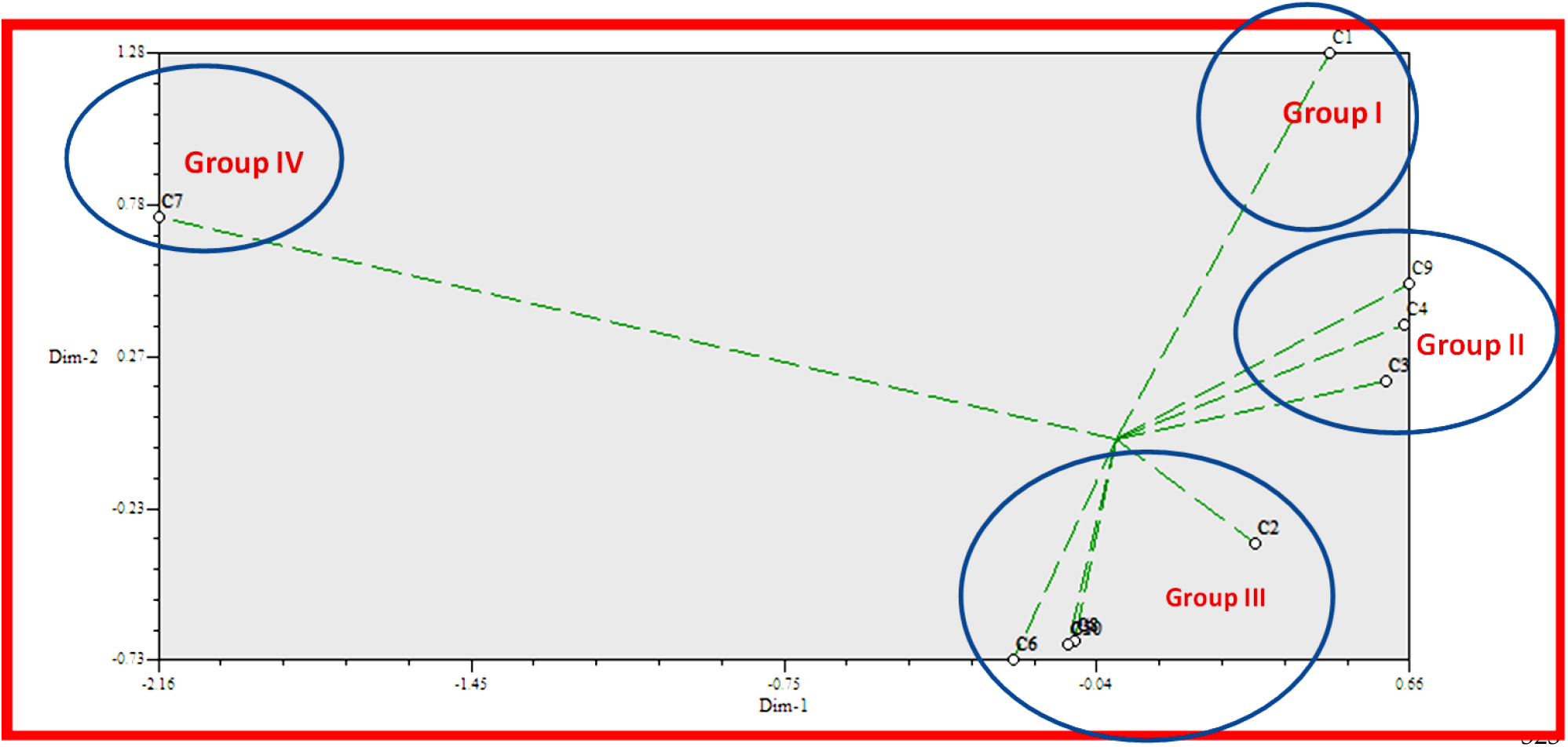
Based on Euclidian distance, the principal component analysis (PCA)-2D graphical association among the indicator plants treated with Parthenium leaf, stem, and flower with methanol extract.

**Figure 5:**
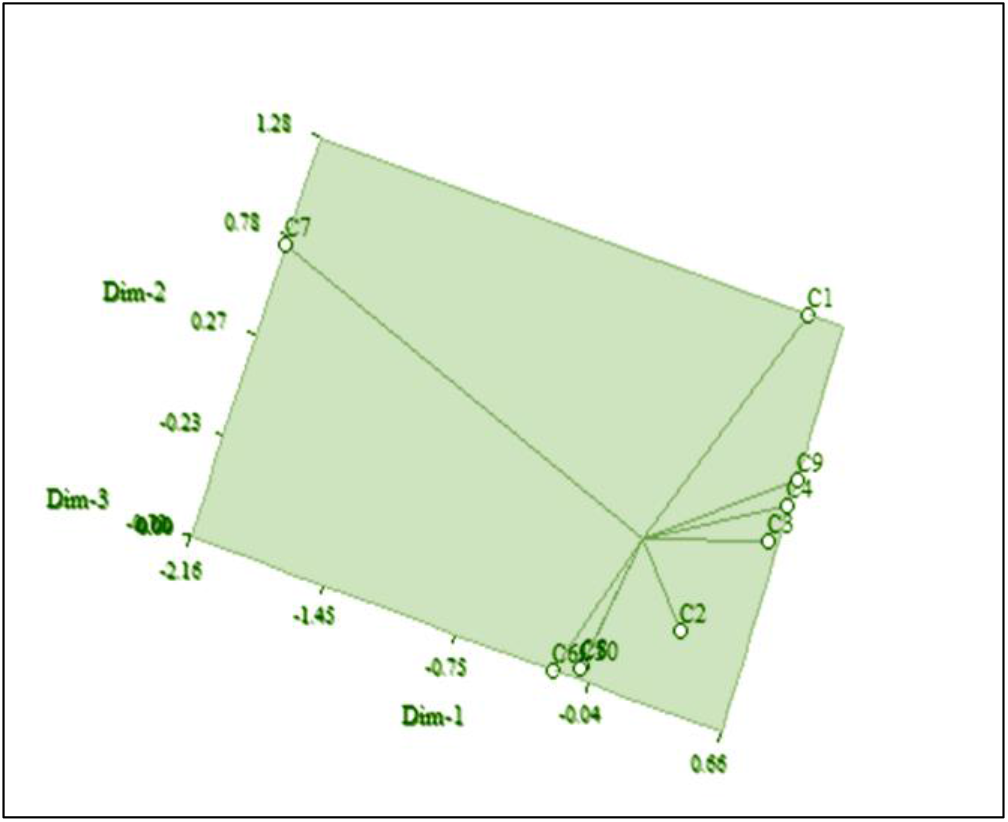
Based on Euclidian distance, principal component analysis (PCA)-3D graphical association between the indicator plants treated with Parthenium leaf, stem, and flower with methanol extract

### 3.4 Identified phenolic derivatives from LC-MS analysis

The identified phenolic derivatives of *P. hysterophorus* plant parts with methanolic extracts through LC-MS analysis are listed in Table 6. The leaf, stem, and flower extracts of *P. hysterophorus* have diverse chemical compositions. A total of 7 Phenolic derivatives were detected from methanol extract of *P. hysterophorus* different parts through LC-MS analysis (Table 6) (Figure 6). These phenolic derivatives are responsible for inhibition to other plants, autotoxic, and dermatitis. Parthenin and other phenolic acids found in the leaf and flower extracts include vanillic acid, caffeic acid, quinic acid, anisic acid, chlorogenic acid, and ferulic acid, contrary Parthenin, vanillic acid found in the stem extract. The amount and kind of chemicals discovered in each plant were found to be proportional to herbicidal action. As a consequence, the compound of the various plant parts inhibited indicator plant germination and seedling growth, with the extraction of leaf having a greater inhibitory influence than the other plant parts.

**Table 6:**
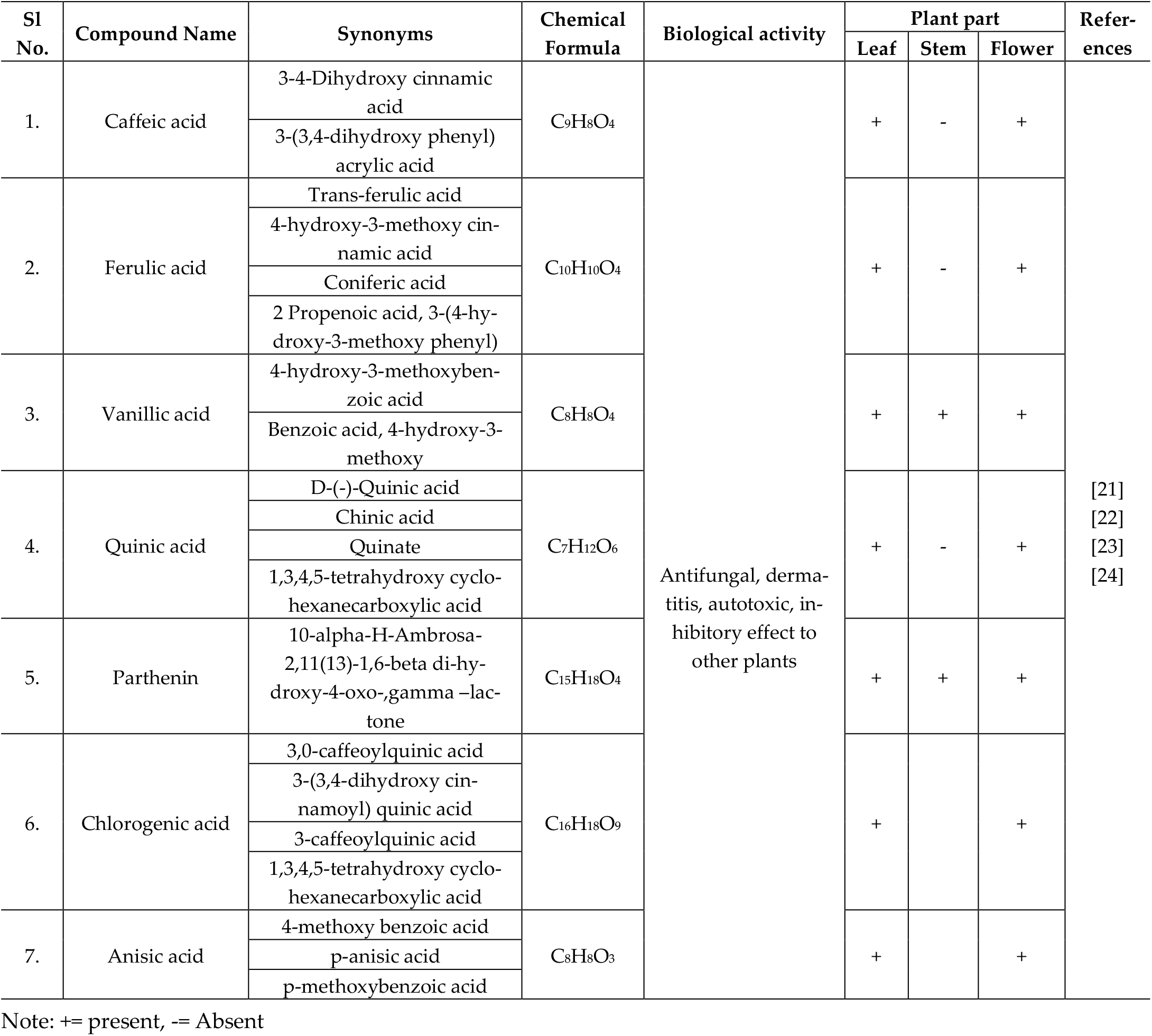
Phenolic derivatives found from methanol extract of *Parthenium hysterophorus* different parts through LC-MS analysis

**Figure 6.**
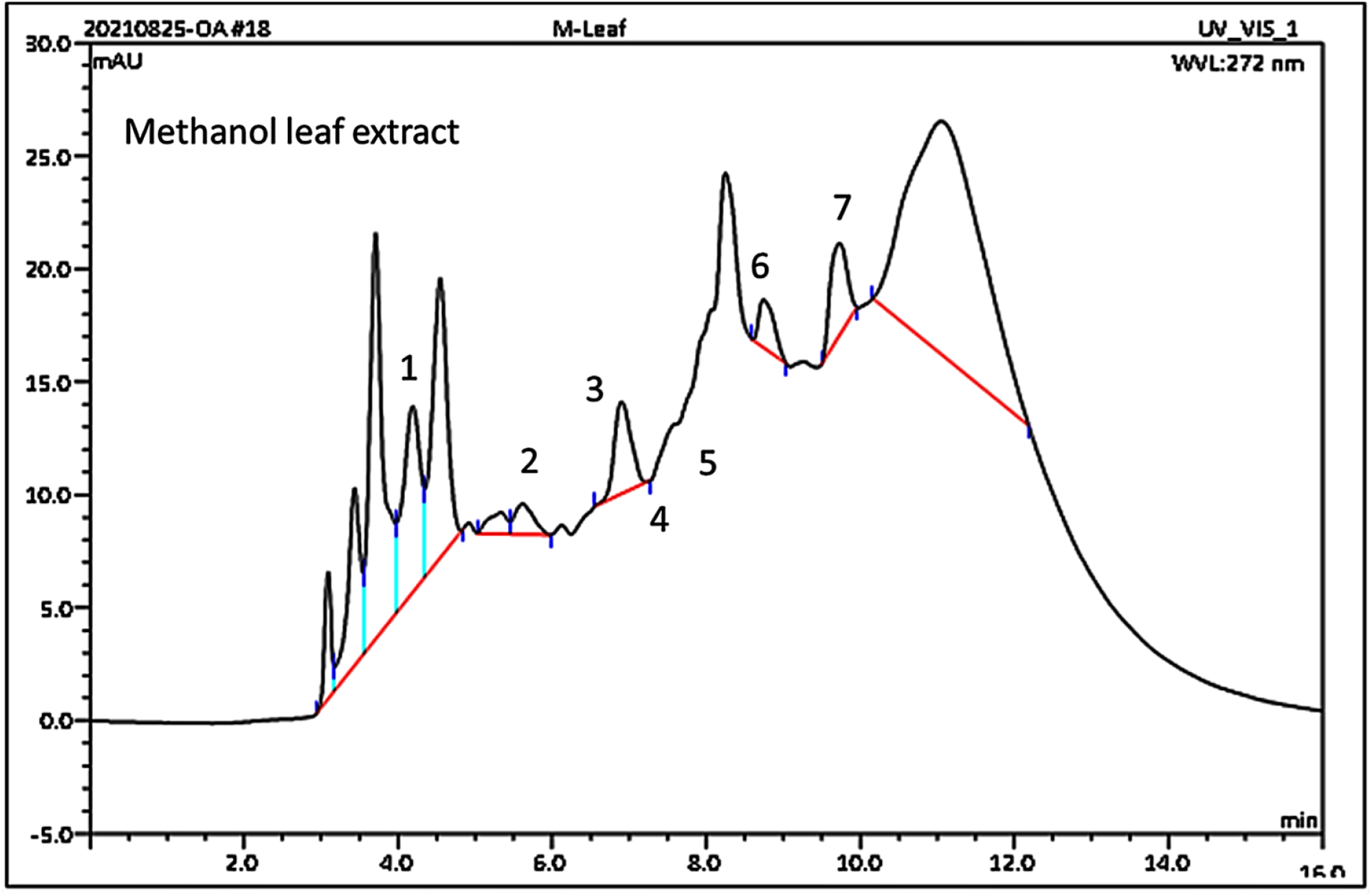
Chromatograms of standard compounds from leaf extract of *P. hysterophorus* (1. Parthenin, 2. Quinic acid, 3. Chlorogenic acid, 4. Vanillic acid, 5. Caffeic acid, 6. Ferulic acid, and 7. Anisic acid)

## 4. Discussion

*P. hysterophorus* with methanol extract influenced germination (%) and growth of seedling of nine different crops (*V. subterranean, R. sativus, C. maxima, C. sativus, S. lycopersicum, C. annum, Z. mays, A. esculentus, D. carota*) and two weed species (*D. sanguinalis* and *E. indica*). In a dose-dependent way, all portions of Parthenium extracts affected germination, radicle length, and hypocotyl length in the tested species. Because of its exceptional strength, efficacy, and consistency in preventing germination and seedling development, extracts of Parthenium leaf were the most promising. Plant extracts are hypothesized to decrease germination through having osmotic potential on the rate of absorption, which in turn affects germination and, in particular, cell elongation [25].

Wheat, maize, and horse gram seedling growth is inhibited by extracts of *P. hysterophorus* methanol extract. Its demonstrated greater inhibitory power, in comparison to the aqueous extract [26]. Dhawan & Gupta [15] reported that the extraction of different active phytochemicals with flavonoid concentration works best using methanol as an extraction solvent. The germination of V. radiata seeds were tested for up to 120 hours using methanol crude Parthenium extracts and it was discovered that there is a considerable difference in germination kinetics between the treatments of methanol crudes. [27].

Tef germination was significantly reduced at intermediate to higher concentrations when Parthenium flower and leaf extracts were used. This suggests that inhibitory compounds are present in larger concentrations in flower and leaf than in stem and root sections [28,29]. The fact that roots came into direct touch with the extract and then with inhibitory compounds, as reported in previous research with a variety of crops and weeds [30,31].

The aerial parts extract of *P. hysterophorus* had a substantial influence on germination of seed, radicle and hypocotyl length reduction in this investigation. These effects grew stronger as the concentration level increased. These discoveries are consistent with those of Mulatu *et al*. [32] and Mersie and Singh [33] who discovered a robust link between greater *P. hysterophorus* aqueous extract concentrations and increased poisonousness to agronomic crops and weeds. The effects of secondary metabolites generated by *P. hysterophorus* aerial parts on growth and development in Bambara groundnut weeds and chosen species. Phytochemicals isolated from *P. hysterophorus* stems, leaves, and flowers methanol extracts were competent to alter crops and weed seedling sprouting and development. Similarly, Motmainna *et al*. [34] discovered that *P. hysterophorus* extract had a considerable impact on the germination and development of the weed species. The degree of inhibition raised when the concentration of the extract was increased. Radicle growth is more vulnerable to allelopathic plant extracts than other organs due to radicles is the first tissue to be shown to phytotoxic chemicals and have a more absorbent tissue than other parts [35,36], and/or the root apical meristem has a low mitotic division rate [37]. Furthermore, allelopathic elements can suppress the production of radicle and epidermis by altering genes involved in cellular characterization [38]. Parthenium extract was more effective than the *B. alata* and *C. rutidosperma* extract [34]. This is in consistent with [39], who discovered that extracts of allelopathic plant have a stronger effect on radicle length than hypocotyl development. This could be due to the roots are the initial to attract allelochemicals substances, from the atmosphere.

The survivability rate of the target plants was inhibited by varying doses of *P. hysterophorus* leaf, stem, and flower methanol extracts. Maximum doses of methanol extracts included more inhibitory chemicals, resulting in more inhibition. In the same way, Han *et al*. [40] had reported that the phytotox-icity of *P. hysterophorus* extracts was concentration-dependent, and phytotoxicity rose as extract concentration was raised. It was also claimed that the leaf extract had a more inhibitory allelopathic activity than other vegetative portions, and phytochemical research had already revealed a larger accumulation of growth inhibitors in *P. hysterophorus* leaves [41]. At all doses examined, the extracts inhibited *P. minor* germination, and when extract concentrations increased then inhibition increased [25]. However, extracts from the leaves had a higher level of toxicity than extracts from the stem [42].

Different plant species’ susceptibility to inhibitory chemicals has been documented for a variety of causes. Msafiri *et al*. [43] observed that both tested species showed substantial allelopathic effects of *P. hysterophorus* seed and leaf aqueous extract on seed sprouting, root and hypocotyl length, fresh and dry mass. According to Kobayasi [44] because of each species’ have biological characteristics. The seed structure and seed coat penetrability can also play a role in different reactions to similar allelopathic extract [45]. Higher concentration reduced the seedling length of all the test crops but, sweet gourd, corn, and cucumber were less sensitive than other crops. This may be due to genotypic variation in response to the higher concentration of extracts. Similar results of inhibitory effect were observed by Aslani *et al*. [42]. These findings agreed with Aslani *et al*. [46], phytotoxic compounds are more vulnerable to smaller plants, he said. These findings matched those of numerous prior research that found that phytotoxin reactions differed by species.

The phytochemical screening revealed a huge number of compounds in the *P. hysterophorus* ex-tracts, some of which have previously been identified as poisons in several investigations [23,47,48]. Furthermore, various plant sections of *P. hysterophorus* contained a different number of compounds. The quantity of toxic compounds was more in the leaf than the other plant parts; as a result, the leaves have a stronger inhibitory effect. *P. hysterophorus* leaves released allelochemicals into the soil by leaching or decomposition, and have the potential to impair the development of other plants by altering the physicochemical properties of soil, according to Dogra & Sood, [49]. Arowosegbe & Afolayan [50] also found that beetroot (*Beta vulgaris* L.), Turnip (*Brassica rapa* L.), and carrot (*D. carota* L.) were all inhibited more by *Aloe ferox* Mill. leaf than by the root extract. The suppressive influence of extracts, according to Verdeguer *et al*. [51] is determined by the extract’s chemical makeup as well as the plant sections to which it is applied. These findings are consistent with those of Javaid and Anjum [52] and Verma *et al*. [53] who discovered that parthenin and other phenolic acids such as caffeic acid, vanillic acid, anisic acid, chlorogenic acid, and para hydroxy benzoic acid are the most responsible for plant growth inhibition.

## 5. Conclusions

The study demonstrated that methanol extracts of *P. hysterophorus* had phytotoxicity on the germination, growth, and development of tested plants. When the concentration of extracts was raised, the rate of germination and seedling growth reduced in comparison to the control, indicating that *P. hysterophorus* was phytotoxic. Moreover, the extracts’ phytotoxic effects were reliant on the target species, concentration of extracts, and plant types. Fifty percent inhibitory concentrations (EC_50_) value of *P. hysterophorus* leaf extract showed more phytotoxic than the stem and flower extract. According to the findings, it was clear that the highly susceptible plants were *Raphanus sativus*, *Solanum lycopersicum*, *Capsicum frutescens, Abelmoschus esculentus, Daucus carota, Digitaria sanguinalis*, and *Eleusine indica*. On the other hand, 7 known phenolic derivatives were identified from the *P. hysterophorus* extract which was responsible for inhibition. Given the hopeful results of *P. hysterophorus* extract, this plant could possibly be studied next in the hopes of developing a herbicide based on natural products for green agriculture that is sustainable. However, in order to offer farmers with useful suggestions, *P. hysterophorus* impacts in actual crop field settings must be validated after bioassay trials. To see if it may be used to develop future alleloherbicides as structural leads, more research on isolation, characterization, and determination of the herbicidal activity of the chemical components in *P. hysterophorus* extract, particularly from the leaf extract, is needed.

## Author Contributions

Conceptualization, A. S. J.; methodology, A. S. J., N. B. A., P. A. and M. S. A. H.; validation, A. S. J., M. K. U., N. B. A., M. S. A. H. and H. M. K. B.; formal analysis, H. M. K. B. and F. R.; investigation, H. M. K. B and F. R.; resources, H. M. K. B. and A. S. J.; data curation, H. M. K. B.; writing—original draft preparation, H. M. K. B.; writing—review and editing, A. S. J., M. K. U., M. S. A. H., P. A., M. A. H. and A.H.; visualization, H. M. K. B and F. R.; supervision, A. S. J., M. S. A. H., M. K. U., and N. B. A.; project administration, A. S. J. and H. M. K. B; All authors have read and agreed to the published version of the manuscript.

## Acknowledgments

Many thanks go to the Ministry of Agriculture (MoA), of the People’s Republic of Bangladesh, Bangladesh Agricultural Research Council (NATP Phase-II Project, BARC), Bangladesh Agricultural Research Institute (BARI) (research grant: vote number 6282506), for providing financial support and Universiti Putra Malaysia (UPM) for assistance. Thankfulness also goes to Mr. Mohd Yunus Abd. Wahab and Mr. Azhar for their assistance in conducting the study.

## Informed Consent Statement

Not applicable

## Data Availability Statement

Not applicable

